# Body size is a better predictor of intra- than interspecific variation of animal stoichiometry across realms

**DOI:** 10.1101/2024.01.22.576743

**Authors:** Mark P. Nessel, Olivier Dézerald, Julian Merder, Karl Andraczek, Ulrich Brose, Michał Filipiak, Michelle Jackson, Malte Jochum, Stan Harpole, Helmut Hillebrand, Shawn J. Leroux, Renske Onstein, George L W Perry, Rachel Paseka, Amanda Rugenski, Judith Sitters, Erik Sperfeld, Maren Striebel, Eugenia Zandona, Hideyuki Doi, Nico Eisenhauer, Vinicius F. Farjalla, Nicholas J. Gotelli, James Hood, Pavel Kratina, Eric K. Moody, Liam N. Nash, Anton M. Potapov, Gustavo Q. Romero, Jean-Marc Roussel, Stefan Scheu, Julia Seeber, Winda Ika Susanti, Alexei Tiunov, Angélica L. González

**Affiliations:** Department of Integrative Biology, University of Guelph, Guelph, ON, Canada; Center for Computational and Integrative Biology, Rutgers University; Camden, 08103, USA; DECOD (Ecosystem Dynamics and Sustainability), INRAE, Institut Agro, IFREMER, 35042 Rennes, France; Department of Global Ecology, Carnegie Institution for Science; Stanford, CA 94305, USA; University Leipzig, Department of Life Sciences, Systematic Botany and Functional Biodiversity, Johannisallee 21, 04103 Leipzig, Germany; EcoNetLab, German Centre for Integrative Biodiversity Research Halle-Jena-Leipzig, Leipzig, Germany; Institute of Biodiversity, Friedrich-Schiller-University Jena, Jena, Germany; Life History Evolution Team, Institute of Environmental Sciences, Faculty of Biology, Jagiellonian University, Gronostajowa 7, 30-387 Kraków, Poland; Department of Biology, University of Oxford, Oxford, UK; Department of Global Change Ecology, Biocenter, University of Würzburg, Würzburg, Germany; German Centre for Integrative Biodiversity Research (iDiv) Halle-Jena-Leipzig, Puschstrasse 4, 04103 Leipzig, Germany; Helmholtz Center for Environmental Research – UFZ, Department of Physiological Diversity, Permoserstrasse 15, 04318 Leipzig, Germany; Institute for Chemistry and Biology of Marine Environment [ICBM], Carl-von-Ossietzky University Oldenburg, Schleusenstrasse 1, 26382 Wilhelmshaven, Germany; Helmholtz-Institute for Functional Marine Biodiversity at the University of Oldenburg [HIFMB], Ammerländer Heerstrasse 231, 26129 Oldenburg; Department of Biology, Memorial University of Newfoundland, St. John’s, NL, Canada, A1C5S7; Naturalis Biodiversity Center, Darwinweg 2, 2333 CR Leiden, the Netherlands; School of Environment, University of Auckland, Auckland, New Zealand; ASRC Federal / NASA Headquarters, 300 E St SW, Washington, DC 20546, USA; Odum School of Ecology, University of Georgia, Athens, Georgia, 30602, USA; Department of Biology, Vrije Universiteit Brussel, Pleinlaan 2, 1050 Brussels, Belgium; B-WARE Research Centre, Nijmegen, the Netherlands; University of Greifswald, Zoological Institute and Museum, Animal Ecology, Loitzer Str. 26, 17489 Greifswald, Germany; Department of Ecology, State University of Rio de Janeiro, Rio de Janeiro, RJ 20550-013, Brazil; Graduate School of Informatics, Kyoto University, Yoshida-honmachi, Sakyo-ku, Kyoto, 606-8501, Japan; Institute of Biology, Leipzig University, Puschstrasse 4, 04103 Leipzig, Germany; Department of Ecology, Federal University of Rio de Janeiro, Rio de Janeiro, RJ 21941-590, Brazil; Department of Biology, University of Vermont, Burlington VT 05405 USA; Aquatic Ecology Laboratory, Department of Evolution, Ecology, and Organismal Behavior, The Ohio State University, 230 Research Center, 1314 Kinnear Road, Columbus, OH, 43212, USA; Translational Data Analytics Institute, The Ohio State University, Columbus, OH, USA; School of Biological and Behavioural Sciences, Queen Mary University of London, Mile End Road, London E1 4NS, UK; Department of Biology, Middlebury College, Middlebury, VT, 05753, USA; Laboratory of Multitrophic Interactions and Biodiversity, Department of Animal Biology, Institute of Biology, University of Campinas (UNICAMP), Campinas, Brazil; Department of Animal Ecology, J.-F. Blumenbach Institute for Zoology and Anthropology, University of Göttingen, Untere Karspüle 2, 37073 Göttingen, Germany; Instiute for Alpine Enviroment, Eurac Research, Drususallee 1, 39100 Bozen/Bolzano, Italy; Universität Innsbruck, Department of Ecology, Technikerstrasse 25, 6020 Innsbruck, Austria; University of Göttingen, J.F. Blumenbach Institute of Zoology and Anthropology, Untere Karspüle 2, 37073 Göttingen, Germany; A.N. Severtsov Institute of Ecology and Evolution, Russian Academy of Sciences, Leninsky prospect 33, 119071, Moscow, Russia; Southern Branch of the Joint Russian-Vietnamese Tropical Research and Technological Center, Ho Chi Minh City, Vietnam; Department of Biology, Rutgers University; Camden, 08103, USA

**Keywords:** Aquatic, Invertebrates, Nitrogen, Phosphorus, Scaling, Terrestrial, Vertebrates

## Abstract

Animal stoichiometry affects fundamental processes ranging from organismal physiology to global element cycles. However, it is unknown whether animal stoichiometry follows predictable scaling relationships with body mass and whether adaptation to life on land or water constrains patterns of elemental allocation. To test both interspecific and intraspecific body-size scaling relationships of the nitrogen (N), phosphorus (P), and N:P content of animals, we used a subset of the StoichLife database encompassing 9,933 individual animals (vertebrates and invertebrates) belonging to 1,543 species spanning 10 orders of magnitude of body size from terrestrial, freshwater, and marine realms. Across species, body mass did not explain much variation in %N and %P composition, although the %P of invertebrates decreased with size. The effects of body size on species elemental content were small in comparison to the effects of taxonomy. Body size was a better predictor of intraspecific than interspecific elemental patterns. Between 42 to 45% in intraspecific stoichiometric variation was explained by body size for 27% of vertebrate species and 35% of invertebrate species. Further, differences between organisms inhabiting aquatic and terrestrial realms were observed only in invertebrate interspecific %N, suggesting that the realm does not play an important role in determining elemental allocation of animals. Based on our analysis of the most comprehensive animal stoichiometry database, we conclude that (i) both body size and realm are relatively weak predictors of animal stoichiometry across taxa, and (ii) body size is a good predictor of intraspecific variation in animal elemental content, which is consistent with tissue-scaling relationships that hold broadly across large groups of animals. This research reveals a lack of general scaling patterns in the elemental content across animals and instead points to a large variation in scaling relationships within and among lineages.

## Introduction

Animal stoichiometry and body size mediate fundamental processes across levels of biological organization from individual-cell metabolism to ecosystem-level functions (Filipiak & Filipiak 2022; Schramski *et al*. 2015; Sterner & Elser 2002). The stoichiometry of life reflects the evolution of a diverse array of life-history strategies related to organismal form and function (Calder 2001; Elser *et al*. 2000a; Sardans *et al*. 2012). Many morphological, physiological, and behavioral traits exhibit predictable scaling relationships with organism size (Brown *et al*. 2000; Enquist *et al*. 2016; Haldane 1926; Peters 1983), and theoretical and empirical work suggests that animal stoichiometry might also be size-dependent (Allen & Gillooly 2009; Elser *et al*. 1996; Gillooly *et al*. 2005; Sterner & Elser 2002). The size scaling of stoichiometry might be similar to that of metabolism, in which a single allometric scaling constant applies over a wide range of species and body sizes (Calder 1984; Kleiber 1932; Savage et al. 2004; West et al. 1997; West & Brown 2005, but see Farrell-Gray & Gotelli 2005; Glazier 2005; Isaac & Carbone 2010). Stoichiometric scaling could have far-reaching implications for ecological and evolutionary research, as anthropogenic activities are driving significant changes in global biogeochemical cycles (Battye *et al*. 2017; Peñuelas *et al*. 2013; Tipping *et al*. 2014), but limited data on stoichiometry and body size across taxa has previously precluded robust tests of these hypothesized stoichiometric relationships.

Size dependency of animal stoichiometry might suggest the existence of fundamental regularities in biological systems (Kempes *et al*. 2019, 2021). Such regularities in biological scaling are viewed as the result of the evolutionary optimization of life-history strategies, which ultimately acts under physical constraints (Kempes *et al*. 2019, 2021; White *et al*. 2022). The elemental content of living organisms and their size-scaling relationships are likely to be governed by an investment in different tissues or biomolecules that are rich in particular chemical elements (Sterner & Elser 2002; Wilder & Jeyasingh 2016). Nitrogen (N) and phosphorus (P) are two such elements found in high concentrations in cells, and are crucial for both physiological (i.e., muscles, vertebrate skeletons and invertebrate exoskeletons) and developmental (i.e., rRNA, proteins) processes in all animals (Extended Data Figure 1). Interspecific scaling relationships describe shifts in biological traits (e.g., metabolism, stoichiometry) as species evolved different body sizes (Glazier 2005). These relationships can be subject to constraints in the evolution (phylogenetic inertia) of these stoichiometric traits (Sterner & Elser 2002). However, in contrast to the plethora of analyses on metabolic scaling, the scaling of animal stoichiometry has only been explored in a few selected invertebrate and fish taxa (Dantas & Attayde 2007; Fagan *et al*. 2002; González *et al*. 2011; Hendrixson *et al*. 2007; Woods *et al*. 2004), often focused on local or regional spatial extents (Allgeier *et al*. 2020; Andrieux *et al*. 2021; Back & King 2013; González *et al*. 2011, 2018; Hambäck *et al*. 2009; Woods *et al*. 2004). Animals, however, display an extraordinary diversity of morphologies and physiologies to withstand the challenges imposed by diverse environments, including terrestrial and aquatic realms, which likely creates diverging constraints on their nutrient demands and body stoichiometry through the distinct selection pressure imposed by gravity and other medium properties (Elser 2006; Elser *et al*. 2000a; Liess *et al*. 2013). Thus, a comprehensive analysis across a large database of animal size and stoichiometry would provide new insights into the critical features governing elemental content.

Gillooly et al. (2005) provided theoretical evidence for the size dependence of whole-body P across species from unicellular eukaryotes to multicellular invertebrates and vertebrates, and 14 orders of magnitude in body size. This relationship supports the Growth Rate Hypothesis (GRH; Elser et al. 2000b), which proposes that small, fast-growing organisms require a large allocation of P-rich rRNA to support growth and biomass production. However, many groups of animals, including most vertebrates and crustaceans, store large amounts of P in structural materials, potentially obscuring the size-scaling of body %P predicted by the GRH (Fabritius *et al*. 2016; Hendrixson *et al*. 2007). Perhaps as a result of P storage, a variety of size-scaling relationships for the elemental content of living organisms, from isometric to allometric, have been described for different groups of plants and animals (Allgeier *et al*. 2020; González *et al*. 2011, 2018; Hendrixson *et al*. 2007; Kerkhoff *et al*. 2005; Woods *et al*. 2004). Similar to metabolism, variation in stoichiometric scaling exponents is partly explained by differences in ontogeny, trophic grouping, and sex (Allgeier *et al*. 2020; Back & King 2013; González *et al*. 2011, 2018; Woods *et al*. 2004). In contrast to microbial life, which may have a common stoichiometric signature (Kempes *et al*. 2019; Neveu *et al*. 2016; Reiners 1986), the structural organization of different types of multicellular organisms may confound the universality of size-scaling of organismal stoichiometry.

Here, we analyze body size-scaling relationships of whole-animal elemental content (i.e., the percentage of the whole-body dry mass made up by a single element in bulk individuals and the molar ratio of elements) using StoichLife, a unique database of invertebrate and vertebrate %N, %P, and N:P body composition (González et al. unpublished).

Interspecific scaling relationships based on species-mean elemental content and body-size scaling should reflect shifts in elemental content with body size, as well as potential phylogenetic inertia. In contrast, intraspecific analyses capture variation over which natural selection can govern evolutionary changes (Stearns 1989). Our overarching hypothesis is that inter- and intraspecific scaling relationships are ultimately explained by allocation of elements to major animal tissues. More specifically, we hypothesize that vertebrates allocate P to bones and N to muscles, while invertebrates allocate P to biomolecules (rRNA; via the GRH) and N to chitinous skeletons and muscles. These animals will also be shaped by the media in which they evolved (i.e., air versus water), leading to predictable relationships between stoichiometry and body size among habitat types. We tested seven specific predictions (Figure 1, Extended Data Figure 1):

**Fig. 1.**
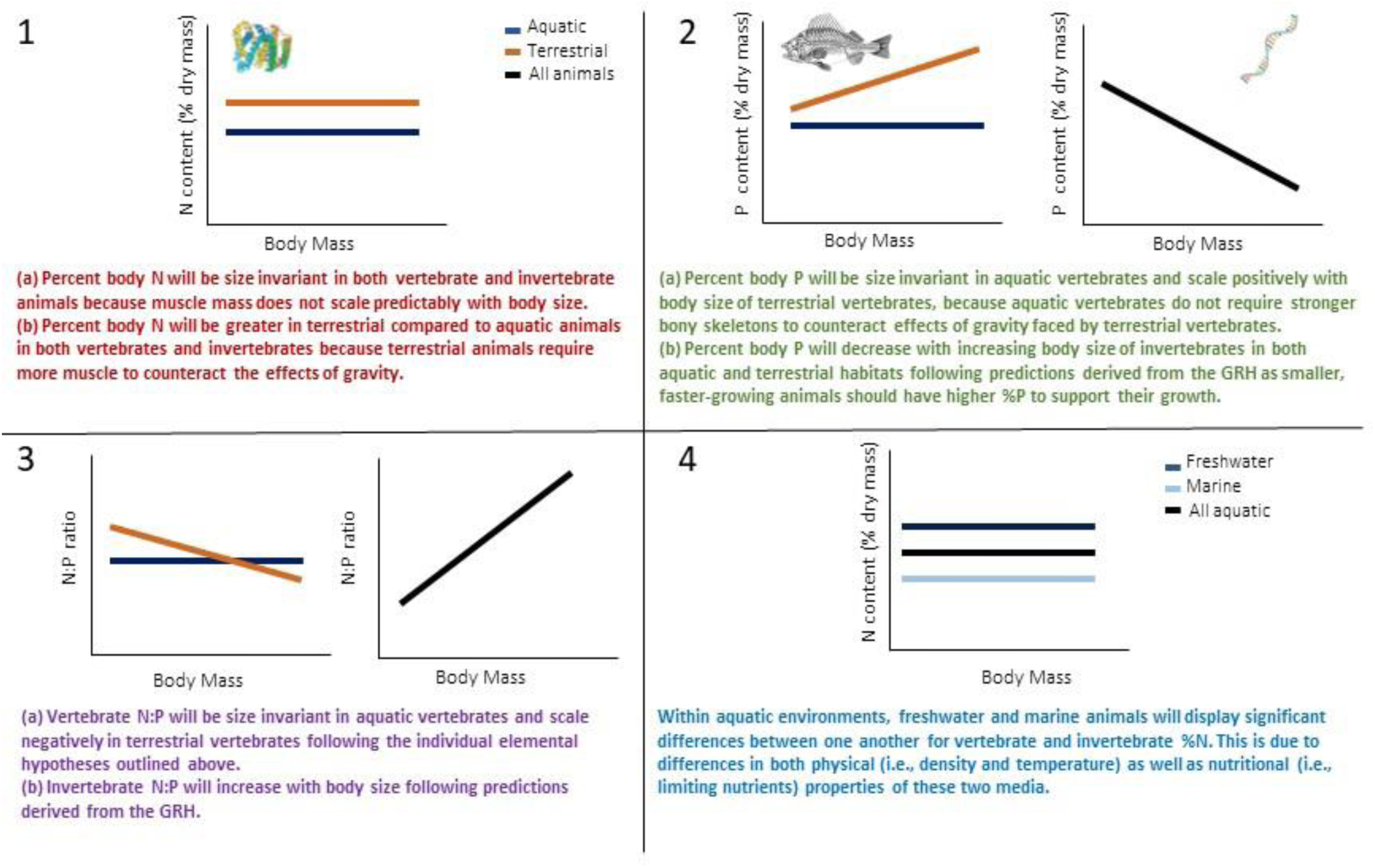
Size scaling predictions and rationale. Predicted size scaling relationships of important body tissues for %N (**1**), %P (**2**), and N:P (**3**), as well as the difference in these scaling relationships in marine and freshwater realms (**4**), and in vertebrates and invertebrates. See Extended Data Figure 1 for complete rationale and evidence for each of these relationships.

(1a) Percent body N will be size invariant in both vertebrate and invertebrate animals because muscle mass does not scale predictably with body size.

(1b) Percent body N will be greater in terrestrial than aquatic animals in both vertebrates and invertebrates because terrestrial animals require more muscle mass to counteract the effects of gravity.

(2a) Percent body P will be size invariant in aquatic vertebrates but scale positively with body size of terrestrial vertebrates, because aquatic vertebrates do not require stronger bony skeletons to counteract the effects of gravity faced by terrestrial vertebrates.

(2b) Percent body P will decrease with increasing body size of invertebrates in both aquatic and terrestrial habitats following predictions derived from the GRH as smaller, faster-growing animals should have higher %P to support their growth.

(3a) Vertebrate N:P will be size invariant in aquatic vertebrates and scale negatively in terrestrial vertebrates following the individual elemental hypotheses (1a and 2a) outlined above.

(3b) Invertebrate N:P will increase with body size following predictions derived from the GRH.

(4) Within aquatic environments, freshwater and marine invertebrates will differ in %N. This is due to differences in physical (i.e., density and temperature), chemical (i.e., salt content, pH, dissolved oxygen), as well as nutritional (i.e., limiting nutrients) properties of these two media. Invertebrate %P follows predictions derived from the GRH, which should hold regardless of the media the organisms inhabit.

## Methods

### Database

We used data from StoichLife, a comprehensive database of organismal (plants and animals) elemental content (%C, %N and %P, and elemental ratios; González et al. unpublished). The StoichLife database is composed of 28,049 individual records. In this study, we only considered animals with %N, %P, N:P, and body mass (dry weight) data available, which resulted in a subset of 9,933 individual and georeferenced records (Extended Data Figure 2). Animals included vertebrates (species = 171; n = 2,861) and invertebrates (species = 1372; n = 7,072) from terrestrial (species = 979; n=3,879), freshwater (species = 516; n = 5,344) and marine (species = 48; n=710) environments. Because some organisms can live in different habitats due to their life cycles, we kept the realm as described in original sources of information (e.g., databases, templates, papers). These species were classified as terrestrial or aquatic according to the place where they mainly feed. For instance, invertebrates with larval stages were classified as aquatic or terrestrial depending on whether the larvae or the winged adult were sampled. This database comprises 10 phyla (Acanthocephala, Annelida, Arthropoda, Chaetognatha, Chordata, Cnidaria, Ctenophora, Mollusca, Nematoda, Platyhelminthes) across 32 classes, 101 orders, and 1,543 (morpho)species. Individual body size in this dataset spans 10 orders of magnitude (ranging from 50 ng to 842 g). Nitrogen content (%N) ranged from 1.13% to 19.54%, phosphorus content (%P) ranged from 0.02% to 8.3%, and N:P ratios ranged from 1.25 to 283. Each element is expressed as the percent of the dry mass in bulk individuals. Elemental ratios (N:P) are expressed as molar ratios.

### Statistical Analyses

First, we investigated interspecific scaling relationships using linear mixed-effects models (LMMs). Prior to this analysis, we logit-transformed percent elemental content (e.g., %N, %P; Warton & Hui 2011), and log_10_-transformed elemental ratios (N:P) and body size. All models for both vertebrate or invertebrate morphospecies (hereafter: species) used mean body size and mean %N, %P, and N:P by species. Regression models included log_10_ body size, realm (i.e., freshwater, marine, terrestrial) as a fixed effect, and their interaction, with logit %N, logit %P or log_10_N:P as response variables. Data on freshwater and marine vertebrate species were pooled, because only three species of marine vertebrates were found in our database. Therefore, prediction (4) was not tested for vertebrate species.

Although phylogenetic relatedness may modulate these elemental-scaling relationships, the extent of our database made it infeasible for us to build a complete, robust phylogenetic tree that encompassed all of the (morpho)species in the subset. Therefore, to account for phylogenetic non-independence, we included “taxonomy” as a random factor in our models, following earlier studies (González *et al*. 2011, 2018; McGill 2008; Tobias *et al*. 2014). In our analyses, we were not able to evaluate a fully-nested taxonomic hierarchy together with our fixed factors due to the lack of model convergence given the complexity of the models. Similarly, we were unable to include both random intercepts and random slopes in our models because of lack of convergence. This lack of convergence is most likely due to too few data points from certain species to allow for calculation of random slopes. Thus, we included a single taxonomic level in our models as a random intercept, which was selected based on the amount of variance explained for a given body element or elemental ratio (i.e., %N, %P, N:P). To do this, we first fitted models with all taxonomic levels independently and selected the taxonomic level with the highest explanatory power as the single random effect in our final model measured as conditional R^2^ value. For vertebrates, taxonomic class explained the most variance in %P and N:P, while family explained most of the variance in vertebrate %N. In invertebrates, phylum explained most of the variance in %N and %P, and family for the N:P ratio (Extended Data Figure 3).

Because body size-scaling relationships of animal elemental content are commonly explored on log-log transformed axes, we also performed analysis for percent elemental (e.g., %N, %P) composition as log_10_-transformed. The outcomes of these analyses were consistent with the logit-transformed results (Extended Data Table 1) further supporting the robustness of our findings. To test for scaling relationships, we fitted separate models for vertebrates and invertebrates. We estimated marginal means via the R package *emmeans* (Lenth 2022) for realm groups, to test the overall and within-realm group effects of body size on elemental content and to compare the slopes and intercepts in each realm. The intercept for each realm is the predicted value at a dry body size of 1 g, which follows from setting the log-body size term in the model to 0. The *P*-values for all model comparisons (i.e., overall effect of body size, effect of body size within each realm, pairwise comparisons between realms) were adjusted using the Benjamini-Hochberg procedure, which controls for the false discovery rate (Pike 2011). To facilitate visualization, %N and %P were back-transformed from logit-transformation (elemental content = e^y^/(1+e^y^)) to median predictions on the original scale.

**Table 1.**
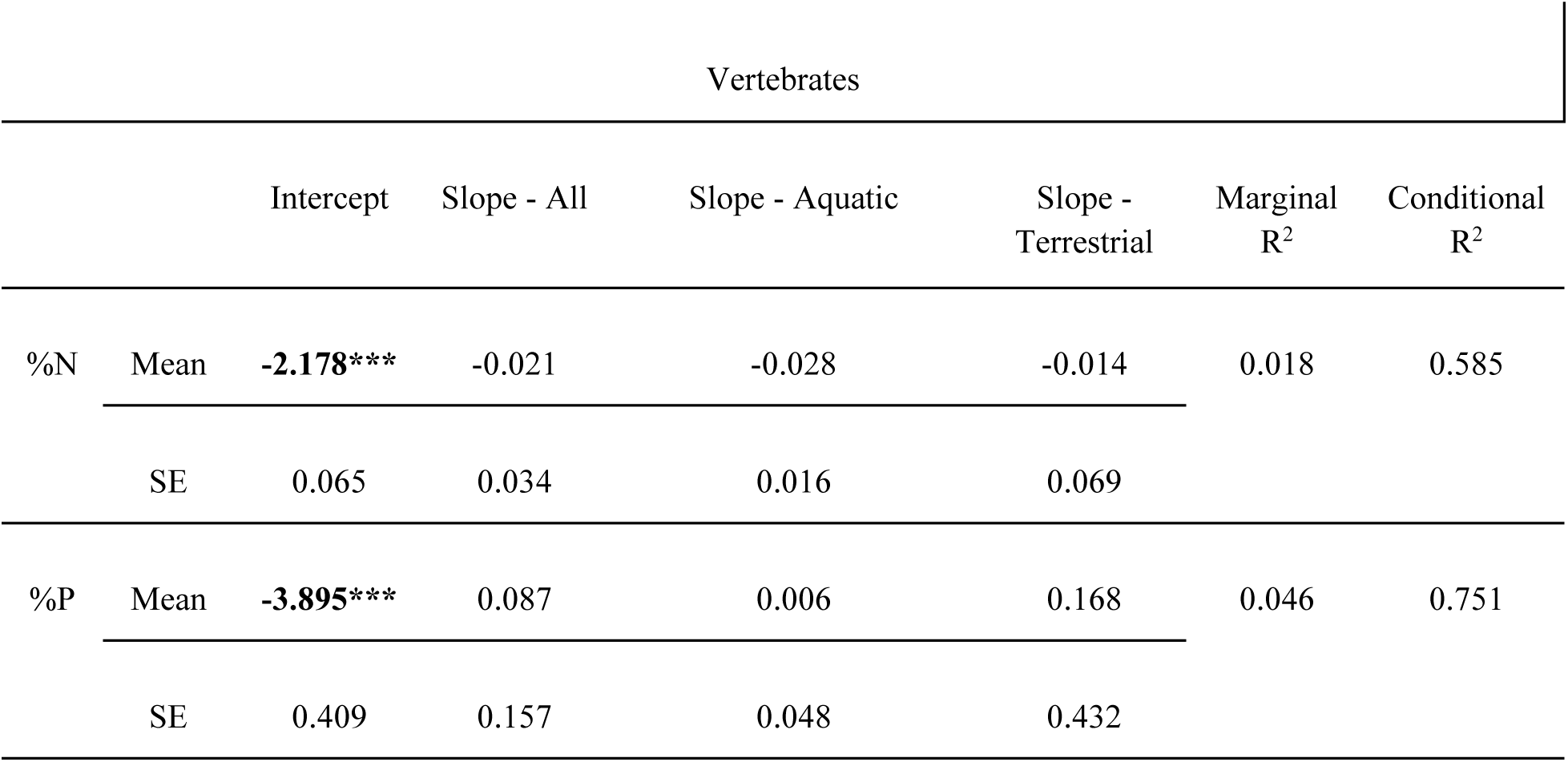
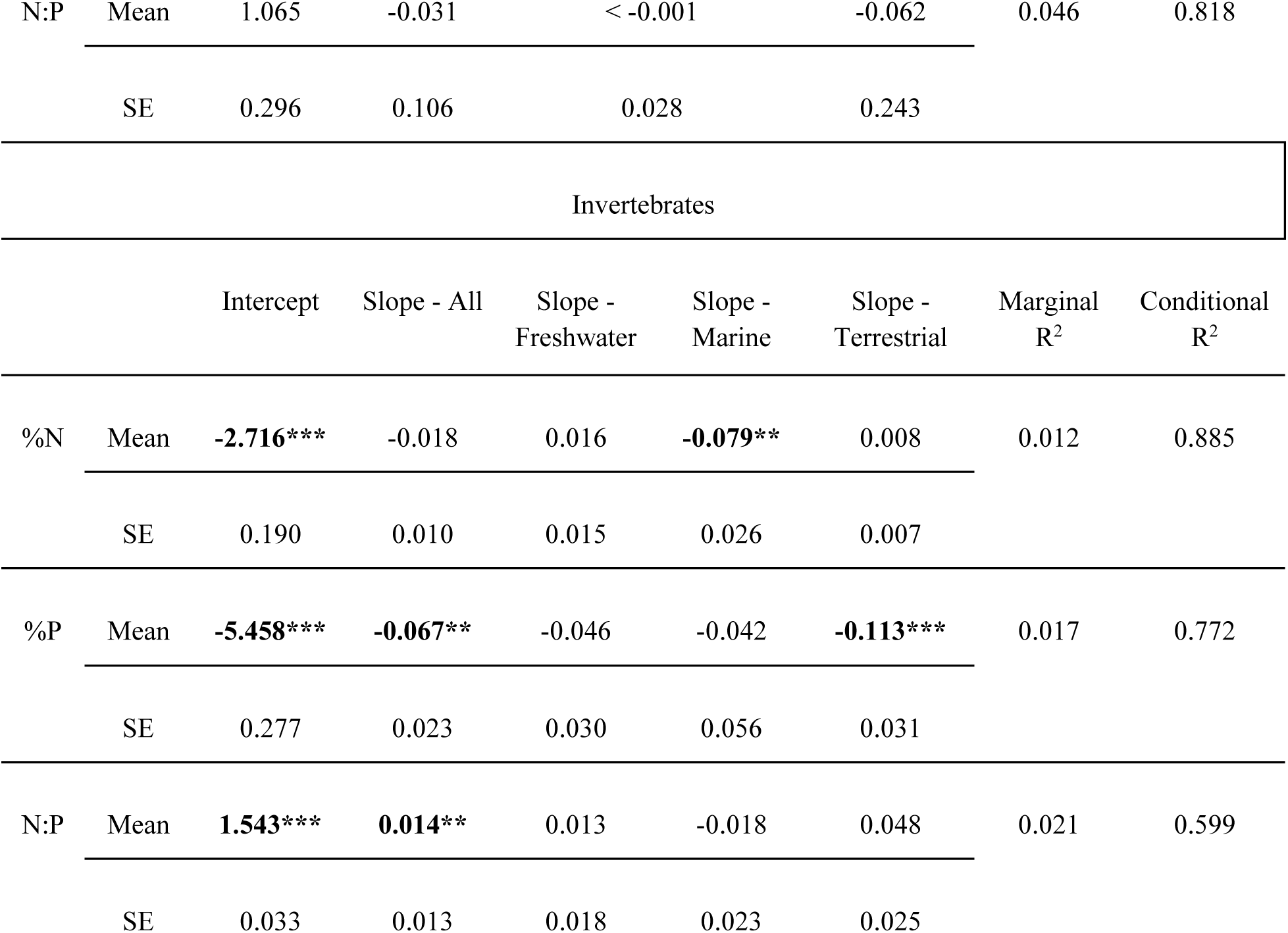
Output of LMMs for %N, %P, and N:P ratios of vertebrates and invertebrates for all animals regardless of taxa (interspecific scaling). The slopes and intercepts of different realms were obtained using estimated marginal means. Predictions are at the population level (fixed effects). The intercept column represents the marginal mean of both groups at a body mass of 1. Significant values in the intercept column indicate the value at body mass 1 is significantly different from 0 when averaged across realms. The column “Slope - All’’ represents the marginal mean slope across realms, so that its magnitude is showing the average effect. The slopes of each realm are also shown. Means and standard errors (SE) are given for all fixed-effect terms as marginal, and fixed plus random effect terms as conditional R^2^ values, for the full model. The N and P relationships are logit-log_10_ transformed, while the N:P ratio relationships are log_10_-log_10_ transformed. Significant values are indicated by bold font and asterisks (*** = *P* <0.001 ** = *P* < 0.01 * = *P* < 0.05).

| Vertebrates |  |  |  |  |  |  |  |
| --- | --- | --- | --- | --- | --- | --- | --- |
| | | Intercept | Slope - All | Slope - Aquatic | Slope - Terrestrial | Marginal $R^2$ | Conditional $R^2$ |
| %N | Mean | **-2.178\*\*\*** | -0.021 | -0.028 | -0.014 | 0.018 | 0.585 |
|  | SE | 0.065 | 0.034 | 0.016 | 0.069 |  |  |
| %P | Mean | **-3.895\*\*\*** | 0.087 | 0.006 | 0.168 | 0.046 | 0.751 |
|  | SE | 0.409 | 0.157 | 0.048 | 0.432 |  |  |

|  |  |  |  |  |  |  |  |  |
| --- | --- | --- | --- | --- | --- | --- | --- | --- |
| N:P | Mean | 1.065 | -0.031 | < -0.001 | -0.062 | 0.046 | 0.818 |  |
|  | SE | 0.296 | 0.106 | 0.028 | 0.243 |  |  |  |
| Invertebrates |  |  |  |  |  |  |  |  |
|  |  | Intercept | Slope - All | Slope -<br>Freshwater | Slope -<br>Marine | Slope -<br>Terrestrial | Marginal<br>R<sup>2</sup> | Conditional<br>R<sup>2</sup> |
| %N | Mean | **-2.716\*\*\*** | -0.018 | 0.016 | **-0.079\*\*** | 0.008 | 0.012 | 0.885 |
|  | SE | 0.190 | 0.010 | 0.015 | 0.026 | 0.007 |  |  |
| %P | Mean | **-5.458\*\*\*** | **-0.067\*\*** | -0.046 | -0.042 | **-0.113\*\*\*** | 0.017 | 0.772 |
|  | SE | 0.277 | 0.023 | 0.030 | 0.056 | 0.031 |  |  |
| N:P | Mean | **1.543\*\*\*** | **0.014\*\*** | 0.013 | -0.018 | 0.048 | 0.021 | 0.599 |
|  | SE | 0.033 | 0.013 | 0.018 | 0.023 | 0.025 |  |  |

For the intraspecific scaling relationships, we used linear models (LMs) and constrained these analyses to only those species represented by at least six individuals (González *et al*. 2017). As with the interspecific models, we logit-transformed percent elemental content and log_10_-transformed elemental ratios before running the analyses. We did not include “realm” as a factor in these models, because each species in our study unambiguously belonged to only one of the realms. These intraspecific models were grouped by realm, and Wilcoxon or Kruskal-Wallis rank sum tests were conducted on the distribution of slopes to determine if there were significant differences between realms. Wilcoxon tests were conducted on groups with only two realms, while Kruskal-Wallis tests were conducted on groups with three realms. Because power increases with sample size, we plotted the relationship between **α**-significance and effect size (i.e., slope) to inspect whether this was affected by sample size (Jenkins & Quintana-Ascencio 2020), but we found no evidence of bias in either *P*-value and slope distributions (Extended Data Figure 4). Moreover, R^2’^s did not vary predictably with sample size, suggesting that they provide an unbiased estimate of the strength of the elemental-size scaling relationships (Extended Data Figure 4).

Because we were interested in the generality and predictability of the body size-scaling relationships, we focused on two model metrics: the coefficient of determination (R^2^) and the *P*-value of our regression coefficients (Price *et al*. 2014). We scored relationships as size invariant if there was a low R^2^ (∼1%; Cohen 1988), or if the slope coefficient was not significantly different from zero (*P* > 0.05), implying that body size has little or no effect on elemental content. All packages and functions in all statistical analyses and figures were implemented and generated using R ver. 4.1.2 (R Core Team 2021).

## Results

### Interspecific Elemental - Body Size Scaling

The body size-scaling relationships of %N, %P, and N:P ratios varied between vertebrates and invertebrates across all realms (Table 1, see Slope-All). In agreement with *prediction 1a* (Figure 1, Extended Data Figure 1), neither vertebrate or invertebrate %N scaled with body size; however, contrary to our *prediction 2a* (Figure 1, Extended Data Figure 1), body size did not have any effect on vertebrate %P. By contrast, invertebrate %P decreased with body size (slope_logit(P)_ = -0.067, SE = 0.023, *P* = 0.004; Table 1), supporting our *prediction 2b* (Figure 1, Extended Data Figure 1). Body size did not explain significant variation in N:P ratios for vertebrates, but significantly increased in invertebrates (Table 1). These results support our *prediction 3a* for both vertebrates and *prediction 3b* for invertebrates (Figure 1, Extended Data Figure 1). The marginal R^2^ values, which are a measure of the variance explained by body size and realm, were much smaller (average marginal R^2^ = 0.026) than the conditional R^2^ (average conditional R^2^ = 0.735), which additionally includes the variance explained by taxonomy as a random effect (Table 1, Figure 2).

**Fig. 2.**
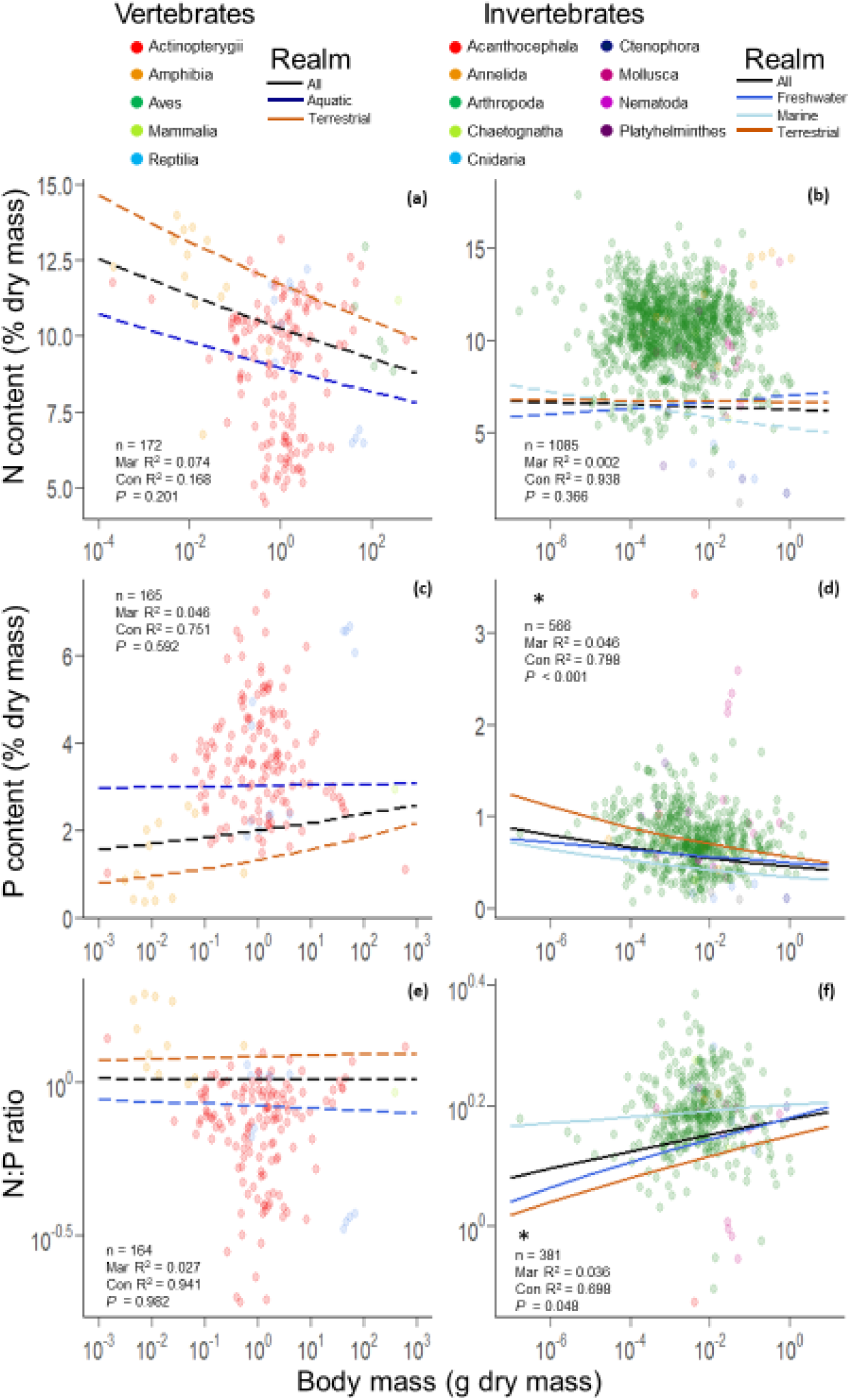
Interspecific elemental scaling relationships. Elemental scaling relationships with body mass for %N (**a, b**), %P (**c, d**), and N:P (**e, f**) in vertebrates (left) and invertebrates (right). Each panel is colored by class (for vertebrates) or phylum (for invertebrates). Black lines indicate the overall effect of body mass on the elemental content of all animals regardless of their group or realm. Colored lines show the effect of body mass for vertebrates (aquatic vs. terrestrial) and invertebrates (freshwater vs. marine vs. terrestrial). These lines do not take into account the random intercepts of each taxonomic group which can cause them to appear off-center from the points. Significant values are shown in bold font and asterisks (*** = *P* <0.001; ** = *P* < 0.01; * = *P* < 0.05). Significant and insignificant relationships are shown using solid and dashed lines, respectively. The R^2^ values are displayed for each model. Elemental content values were back-transformed to median values from logit-transformed values in order to ease interpretation. The R^2^, *P*-values for the overall model, and the number of species (n) for each model are displayed in the panels.

These results showed that both vertebrates and invertebrates from terrestrial realms were similar in %N to those from aquatic realms at a body size of 1 g, which did not support our *prediction 1b* (Table 2, Figure 2a-b). In addition, we found that terrestrial and aquatic vertebrates did not differ in their %N scaling slopes. By contrast, the invertebrate %N scaling slopes from marine, terrestrial, and freshwater realms were statistically different and intersected at small body sizes (Figure 2b). As a result, small marine invertebrates (with body sizes below 10 µg dry mass) tended to have higher %N than small freshwater and terrestrial invertebrates, whereas marine %N was significantly lower at a body size of 1 g (Figure 2b, Tables 1, 2).

**Table 2.** Pairwise comparison between the slopes and intercepts of animals from different realms using estimated marginal means. Intercept comparisons were done at log_10_. Significant values are shown in bold font and asterisks (*** = *P* <0.001; ** = *P* < 0.01; * = *P* < 0.05).

| Slope |  |  |  |  |  |
| --- | --- | --- | --- | --- | --- |
| Vertebrate |  |  |  |  |  |
|  | contrast | estimate | SE | df | *P*-value |
| %N | aquatic - terrestrial | -0.014 | 0.071 | 71.34 | 0.844 |
| %P | aquatic - terrestrial | -0.161 | 0.437 | 17.69 | 0.715 |
| N:P | aquatic - terrestrial | 0.061 | 0.246 | 7.45 | 0.808 |
| Invertebrate |  |  |  |  |  |
|  | contrast | estimate | SE | df | *P*-value |
| %N | **freshwater - marine\*\*** | **0.096** | **0.030** | **1086.38** | **0.004** |
|  | freshwater - terrestrial | 0.007 | 0.017 | 1080.34 | 1.000 |
|  | **marine - terrestrial\*\*** | **-0.088** | **0.027** | **1088.08** | **0.003** |
| %P | freshwater - marine | -0.003 | 0.063 | 569.99 | 1.000 |
|  | freshwater - terrestrial | 0.067 | 0.043 | 561.50 | 0.374 |
|  | marine - terrestrial | 0.071 | 0.064 | 570.01 | 0.812 |
| N:P | freshwater - marine | 0.032 | 0.027 | 359.24 | 0.724 |
|  | freshwater - terrestrial | -0.035 | 0.030 | 364.97 | 0.754 |
|  | marine - terrestrial | -0.067 | 0.034 | 361.43 | 0.146 |
| Intercept |  |  |  |  |  |
| Vertebrate |  |  |  |  |  |

|  | contrast | estimate | SE | df | P-value |
| --- | --- | --- | --- | --- | --- |
| %N | aquatic - terrestrial | -0.059 | 0.135 | 52.07 | 0.661 |
| %P | **aquatic - terrestrial\*\*** | **0.840** | **0.277** | **170.65** | **0.002** |
| N:P | **aquatic - terrestrial\*\*\*** | **-0.580** | **0.166** | **155.34** | **< 0.001** |
| Invertebrate |  |  |  |  |  |
| %N | **freshwater - marine\*\*\*** | **0.364** | **0.081** | **1090.62** | **< 0.001** |
|  | freshwater - terrestrial | -0.092 | 0.046 | 1080.47 | 0.140 |
|  | **marine - terrestrial\*\*\*** | **-0.457** | **0.077** | **1091.12** | **< 0.001** |
| %P | freshwater - marine | -0.064 | 0.175 | 572.15 | 1.000 |
|  | freshwater - terrestrial | -0.063 | 0.112 | 561.33 | 1.000 |
|  | marine - terrestrial | < 0.001 | 0.175 | 572.08 | 1.000 |
| N:P | freshwater - marine | 0.132 | 0.079 | 293.95 | 0.289 |
|  | freshwater - terrestrial | -0.032 | 0.074 | 339.49 | 1.000 |
|  | marine - terrestrial | -0.164 | 0.086 | 278.15 | 0.176 |

In line with *prediction 2a*, we did find that aquatic invertebrate %P was size invariant, however, %P terrestrial vertebrates did not scale positively with body size as we predicted. Invertebrate %P was not significantly different between realms, which support our *prediction 2b* (Table 1, Figure 2d). Further, for vertebrates, the slope of the body size-N:P scaling relationship did not differ significantly between terrestrial and aquatic realms (Table 1, Figure 2e). However, the intercept for vertebrate N:P was significantly higher in terrestrial than aquatic animals (Table 2). Aquatic and terrestrial invertebrates did not differ in their scaling slopes or intercepts for N:P (Table 1, Figure 2f). Unlike %N, we found no significant differences between freshwater and marine invertebrates in %P or the N:P ratio, which only partially supported our *prediction 4* (Table 2, Figure 2). We were only able to test for differences in elemental content and regression coefficients for invertebrates, but not vertebrates, within the aquatic realm (i.e., freshwater and marine) due to the lack of marine vertebrates in our database.

### Intraspecific Elemental - Body Size-Scaling for Vertebrates and Invertebrates

Body size-scaling relationships varied widely among species (Figures 3, 4; Extended Data Table 2). Although only 22% of the vertebrate species (n = 101) and 32% of the invertebrate species (n = 235) showed significant body size relationships for %N, among these significant relationships, body size explained on average 40% (SD 21%) and 43% (SD 24%) of the variance in logit %N, for vertebrates and invertebrates, respectively (Extended Data Table 2). Further, 30% of vertebrate species (n = 99) and 44% of invertebrate species (n = 114) displayed significant scaling relationships for %P, and body size explained 42% (SD 15%) and 47% (SD 22%) of the variance in logit %P in these significant vertebrate and invertebrate relationships, respectively. Finally, body size explained on average 39% (SD 19%) and 49% (SD 24%) of the variance in vertebrate and invertebrate log10 N:P, respectively, but only 26% of vertebrate species (n = 99) and 32% of invertebrate species (n = 93) displayed significant N:P scaling relationships (Extended Data Table 2).

**Fig. 3.**
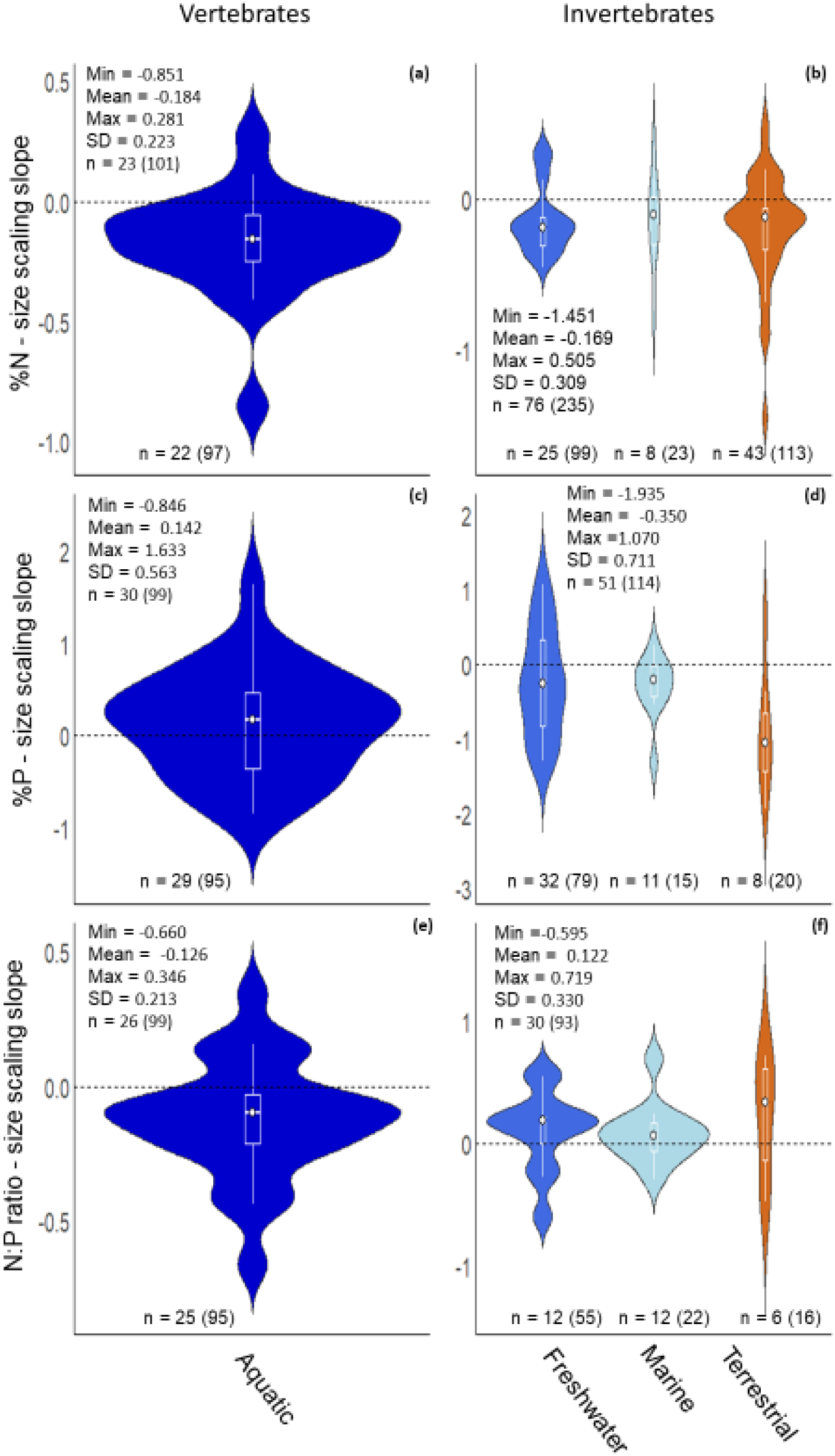
Intraspecific slope distributions. Distribution of slopes estimated for each species from the intraspecific models for %N (**a, b**), %P (**c, d**), and N:P ratios (**e, f**) in vertebrates (left) and invertebrates (right) separated by realm. For each model, elemental content (i.e., %N, %P) was logit-transformed, while N:P ratios were log_10_-transformed. Models included a variable number of species for each group. Only significant relationships (*P* < 0.05) are shown. Relationships were grouped by realm after the analysis. Only aquatic species are shown for vertebrates as there was only one marine species with a significant relationship, so the two groups were combined. The number of species with significant relationships are shown in the violin plots. Due to the limited number of terrestrial vertebrate species with significant relationships only aquatic invertebrate slopes are examined in the violin plots. The white dot represents the median value of the intercept. The panels also hold the ranges, mean, SD of the slopes for each group, the total number of species with significant relationships based on the regression models, and the total number of species tested in parentheses.

Similar to interspecific scaling, we did not find any evidence for the predicted differences between realms for intraspecific body size-scaling relationships. However, we did not have enough marine vertebrate species, so for this test we only compared slopes between freshwater and terrestrial vertebrates (Figure 3). Vertebrate %N ([Wilcoxon] W = 201, *P* = 0.91), %P ([Wilcoxon] W = 195, *P* = 0.94), and N:P ([Wilcoxon] W = 225, *P* = 0.54) and invertebrate %N ([Kruskal-Wallis] chi-squared = 3.32, df = 2, *P* = 0.19), %P ([Kruskal-Wallis] chi-squared = 8.85, df = 2, *P* = 0.012), and N:P ([Kruskal-Wallis] chi-squared = 1.72, df = 2, *P* = 0.42) did not differ significantly between freshwater and terrestrial realms (Figure 3).

## Discussion

Organismal stoichiometry is central to ecology and evolution, but the potential links between stoichiometry and body size have not previously been tested across a large diversity of animals. Based on a novel database, our results demonstrate that size alone explains a substantial fraction of the variance in intraspecific elemental content, but it is a poor predictor of interspecific elemental content. Our results revealed that the %N of marine invertebrates differed significantly from that of freshwater and terrestrial invertebrates, with small marine invertebrates tending to have higher %N than small freshwater and terrestrial invertebrates, and larger marine invertebrates having lower %N then other larger invertebrates. Contrary to theoretical predictions, aquatic vertebrates had higher %P than terrestrial vertebrates at an equivalent body size. Thus, major, nutrient-rich tissues and biomolecules present in all animal bodies (i.e., vertebrate skeletons, muscles) do not seem to be predictable drivers of organismal elemental content (Sterner & Elser 2002). Accounting for taxonomy to control for the large morphological and physiological diversity of animals included in our analyses, allowed us to explain considerably more variance in these scaling relationships. Contrary to predictions by ecological stoichiometry and metabolic theory, our analyses of a global dataset demonstrate that body mass alone is a poor predictor of animal elemental content.

### Inter- and Intraspecific Elemental - Body Size Scaling

We expected vertebrate %N-body size scaling to be driven by protein content (Sterner & Elser 2002; Striebel *et al*. 2017). Extensive empirical work has shown that muscle and skin are major pools of protein in vertebrates (Calder 1984), with muscle contributing 10% to > 60% of body mass (Johnston *et al*. 2011; Muchlinski *et al*. 2012; Schreck *et al*. 2016). These studies have revealed isometric relationships between protein mass and body size for birds (Bennett 1996) and fishes (Breck 2014; Li *et al*. 2016), the latter representing the largest group of vertebrates in our database (71% of all vertebrates). The size-invariance of vertebrate %N supported *prediction 1a* (Table 1, Figure 2a), and confirmed previous work examining vertebrate muscle content scaling (Bennett 1996; Muchlinski *et al*. 2012; Pollock & Shadwick 1994). Likewise, the mean %N slope across vertebrate species did not significantly differ from zero (Figure 3a, Figure 4a). While we did not directly test the scaling of muscle content, %N was size invariant for vertebrates (Table 1, Figure 2a), supporting the idea that the scaling of %N in vertebrates can be explained by muscle investment.

**Fig. 4.**
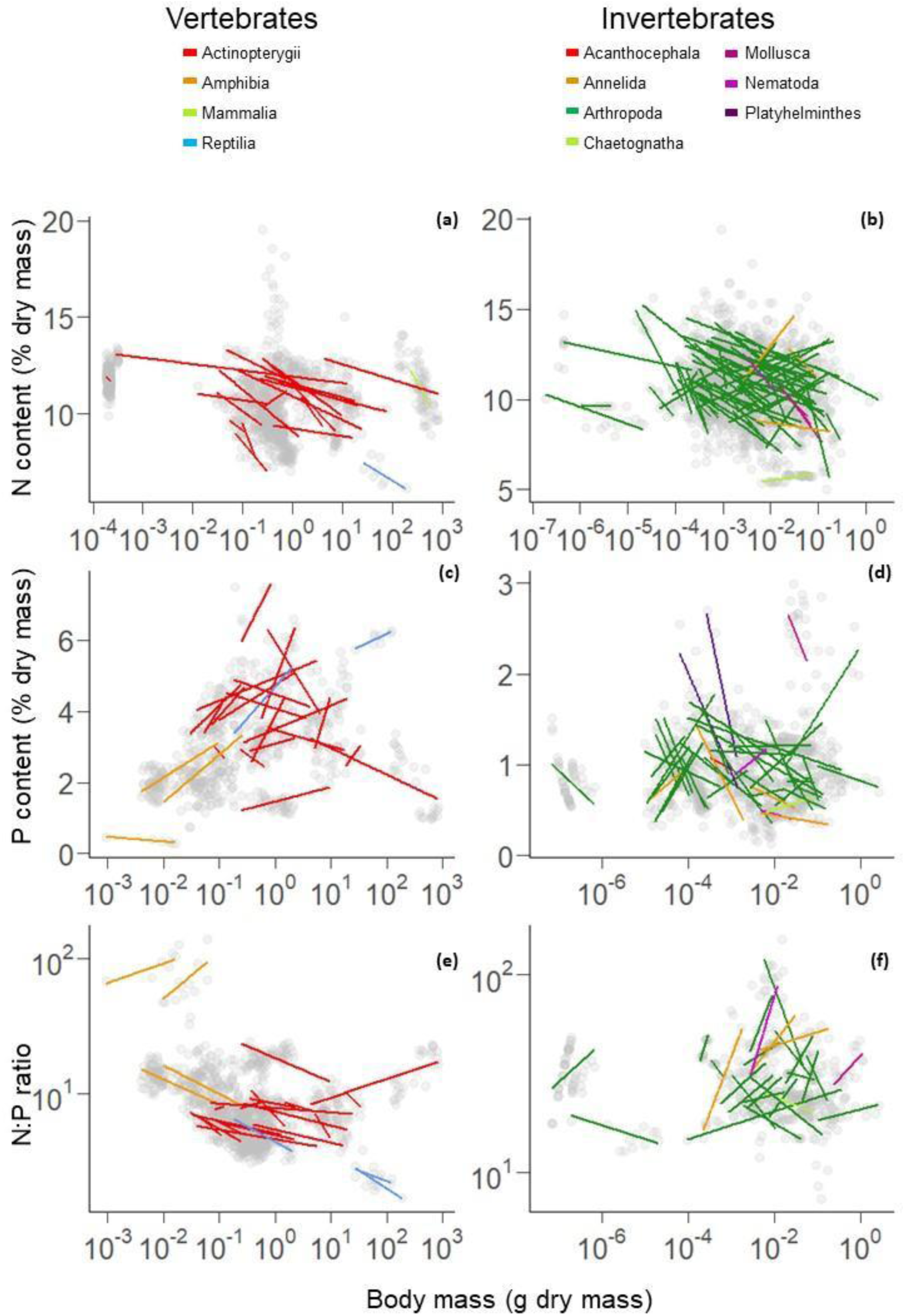
Intraspecific elemental scaling relationships. Intraspecific scaling relationships for %N (**a, b**), %P (**c, d**), and N:P ratios (**e, f**) in vertebrates (left) and invertebrates (right). Each panel is colored by class (vertebrates) or phylum (invertebrates). We used class for vertebrates because they represent a single phylum (i.e., Chordata). Only species with significant relationships (*P* < 0.05) are shown. Elemental content values were back-transformed to median values from logit-transformed values in order to ease interpretation.

The major invertebrate protein pools are in muscles and the exoskeleton of arthropods (96% of invertebrates in our database) (Finke 2007). Indeed, protein represents up to 50% of arthropod exoskeleton mass (Sterner & Elser 2002; Wilder & Barnes 2023). Exoskeletal investment varies from 2% to 80% of the body mass of an organism (Anderson *et al*. 1979; Chapman 2013; Jonas-Levi & Martinez 2017), and can scale both isometrically or allometrically with body size (Glazier *et al*. 2011; Polilov & Makarova 2017; Schramm *et al*. 2018; Whitman 2008). In addition, flight muscle in insects accounts for between 0% and 65% of their body size, across flightless and winged species (Marden 1989, 2000). Insect flight muscles scale at different rates across several insect orders (Marden 1987; Polilov & Makarova 2017), with individual orders ranging from negative allometry (slope = 0.73) to positive allometry (slope = 1.22) (Brown *et al*. 2017; Marden 1987). In our study, the N content of all invertebrate species did not significantly change with body size (Table 1, Figure 2b). We suggest that differences in morphology and physiology within this diverse group of organisms likely lead to broad divergences in the allocation to and scaling of exoskeletal and flight tissues, which could, in part, explain the large stoichiometric variation and the importance of taxonomy underlying our observed scaling relationships (Figure 2b). This explanation is also consistent with the large intraspecific variation observed in our scaling relationships (Figure 3b, Figure 4b). Hence, body size alone is insufficient to explain how animals allocate these protein-rich tissues to build their bodies. Similar to the role of life histories in the scaling of metabolism (Glazier 2005; Jerde *et al*. 2019; Sibly & Brown 2007; White *et al*. 2022), additional information on other traits related to ontogeny and physiology may improve predictions of the scaling of invertebrate N composition.

Skeleton mass is the main driver of whole-body phosphorus in vertebrates, which scales allometrically with a slope between 1.07 and 1.20 in terrestrial vertebrates (Christiansen 2002; Martin-Silverstone *et al*. 2015; Prange *et al*. 1979) and isometrically in aquatic vertebrates (Berrios-Lopez *et al*. 1996; White & Seymour 2011). This size scaling has been explained by an increase in skeletal investment in terrestrial environments to counteract the effects of gravity (Berrios-Lopez *et al*. 1996; White & Seymour 2011). Across vertebrate species, %P was size invariant for terrestrial and aquatic vertebrates (Table 1, Figure 2c), and the slope was not different from zero (Figure 3c). Previous studies examining scaling relationships in aquatic vertebrates have reported %P to be size invariant across multiple species (Dantas & Attayde 2007; Hendrixson *et al*. 2007; Sterner & George 2000). Some studies, however, have reported large differences in %P among vertebrate species, as well as diverse intraspecific scaling relationships (El-Sabaawi *et al*. 2012; Tanner *et al*. 2000). The slopes of %P for individual species in our study ranged from positive (slope = 0.685) to negative (slope = -0.295) (Figures 3c, 4c), suggesting that the variety of skeletal frames (i.e., shape and thickness of bones and scales) could partially explain the observed variation in scaling exponents (El-Sabaawi *et al*. 2016; Hendrixson *et al*. 2007). The lack of a relationship across terrestrial vertebrates in our study should be taken with caution, as this finding is supported by an analysis with only nine species, limiting our inferences on their scaling relationships.

The decrease in invertebrate %P with body size (Table 1, Figure 2d) is consistent with predictions derived from the GRH, which posits that an increase in P-rich RNA content is associated with an increase in %P in small, fast-growing organisms. While the average slope across species was negative, there was substantial variation among species, with some displaying steep positive slopes (Figures 3d, 4d). For very small invertebrates, rRNA can constitute up to 15% of total body weight (Sterner & Elser 2002), representing up to 80% of total body P (Vrede 1999) suggesting that RNA can be a significant driver of whole-body elemental content. Previous paradigms held that non-rRNA-P, including pools of phospholipids and ATP, does not vary with body size in larger animals (Allen & Gillooly 2009; Elser *et al*. 2003; Gillooly *et al*. 2005). However, recent studies have reported large variation in these pools for non-growth-related functions such as structure (e.g., phospholipids) and storage as inorganic polyphosphate (Fabritius *et al*. 2016; He & Wang 2020; Isanta-Navarro *et al*. 2022). In addition, investment in certain growth-related products like cocoons, which are required by many holometabolous insects to complete their metamorphosis, can represent a large amount of P allocation (Filipiak *et al*. 2021). Further, growth limitation by nutrients other than P or energy has been shown to affect relationships between growth, size, and P content (Isanta-Navarro *et al*. 2022; Lukas *et al*. 2011). These mechanisms could lead to a decoupling of body P content and growth, resulting in a much lower proportion of variation in %P being explained by body size and realm than initially predicted by the GRH (González *et al*. 2018; Isanta-Navarro *et al*. 2022).

The N:P ratios of vertebrates were size invariant and increased significantly with body size for invertebrates (Table 1, Figure 2), which is consistent with our prediction of the N:P scaling. These results agree with a lack of relationship between the body size of vertebrates and their N:P found in previous studies (Dantas & Attayde 2007; Hendrixson *et al*. 2007). A potential explanation for the lack of relationship between N:P and body size is that aquatic vertebrates do not increase their skeletal mass proportionally with body size, but instead invest in different body structures (bones and scales), which vary widely in size and density among species independent of changes in body size (Durston & El-Sabaawi 2017; El-Sabaawi *et al*. 2016). We predicted that the N:P ratio of invertebrates would scale positively with their body size. Our results agree with earlier studies showing significant increases in invertebrate N:P ratio with body size (Allgeier *et al*. 2020; Back & King 2013; González *et al*. 2011). We aimed to identify general scaling trends regardless of taxonomic identity. However, we acknowledge that variation in evolutionary history among lineages (e.g., crustaceans vs. insects) may lead to deviations from general patterns and should be investigated further. Recent evidence has also shown that developmental stage can have strong influence on animal stoichiometry (e.g., insects (Fagan *et al*. 2002; Filipiak & Weiner 2017), vertebrates (May & El-Sabaawi 2022)) due to growth-dependent P allocation to RNA and skeletons (Isanta-Navarro *et al*. 2022). We were not able to separate organisms by developmental stage, as this information was not reported for many of the records in our database. Future studies should incorporate evolutionary history, physiological traits and developmental stages in order to quantify how these characteristics drive organismal stoichiometry.

### The Effect of Realm on Size-Scaling Relationships

Contrary to our predictions, neither terrestrial vertebrates or invertebrates had significantly higher %N than aquatic animals (Figure 2, Table 2). Further, both vertebrate and invertebrate species in different realms did not significantly differ in scaling slopes (Figure 3). We predicted a higher %N for terrestrial animals because they are expected to invest more in muscular tissue relative to aquatic animals to support the denser skeletons required to prevent water loss and to move against gravitational forces. Differences between realms in terms of structural allocation have been previously demonstrated in vertebrate muscle content (Denny 1990; Schmidt-Nielsen 1972) and in the cuticle content of some arthropods (Taylor 2018). However, to our knowledge, ours is the first study to specifically test for differences in the %N-size scaling of aquatic and terrestrial animals. The exact mechanisms driving these scaling differences are unclear and require more-explicit analyses of both tissue and elemental content as they relate to body size. We suggest that lower scaling intercepts for aquatic organisms could reflect constraints on the body plan for mechanical adaptations to life in water, such as increases in vertebrate buoyancy (Mansuit & Herrel 2021) and invertebrate exoskeleton thickness (Taylor 2018). Alternatively, diversity in locomotion modes, particularly in aquatic organisms (e.g., drag-based, lifted-based, jetting) may influence elemental-size scaling relationships, if these strategies generate trade-offs between muscle allocation to support increases in body size versus locomotion (Clemente & Richards 2013; Vogel 2008).

Contrary to our expectation, aquatic vertebrates had significantly higher %P than terrestrial vertebrates (Figure 2c, Table 2). Studies explicitly considering P content of terrestrial vertebrates have reported positive scaling relationships with body size (Milanovich *et al*. 2015), in contrast to the size-invariant relationships observed in aquatic vertebrates (Berrios-Lopez *et al*. 1996; Dantas & Attayde 2007; Hendrixson *et al*. 2007). Similar to our expectations for %N, we expected higher %P in terrestrial compared to aquatic vertebrates due to their increased requirements for skeletal investment in a gravitational environment compared to a buoyant environment (Berrios-Lopez *et al*. 1996; White & Seymour 2011). Our results suggest that, despite living in different media, evolutionary pressures on material (bone) investment, and therefore P content, may be acting on terrestrial and aquatic vertebrates under fundamental physical constraints (Kempes *et al*. 2019). Comparisons of bone density in vertebrates inhabiting different realms have found greater bone density in terrestrial than aquatic vertebrates (Fish 2000; Reynolds 1977; Wall 1983). Differences in bone density could translate to variable body %P. However, none of those earlier studies directly compared %P in the body of aquatic and terrestrial vertebrates. Although the specific mechanism behind the lower P investment in terrestrial compared to aquatic organisms is currently unknown, we suggest that structures other than the skeleton might contribute to the body %P in vertebrates affecting body size scaling-relationships. For example, dermal elements in the form of scales and teeth represent important components of P allocation and storage in both groups of vertebrates (Doherty *et al*. 2015), with some fish taxa such as Loricariids allocating large amounts of P to armor-like bony plates (De Andrade Santos *et al*. 2016). However, the limited data describing %P and N:P ratios of terrestrial vertebrates (8% of vertebrate species), limit our inferences on their elemental content-body size scaling. In our analysis, we found no support for terrestrial vertebrates having higher %P than aquatic vertebrates. However, there is a shortage of data on elemental content of terrestrial vertebrates, especially those that are larger than 1 kg dry mass, currently limiting a comprehensive understanding of differences between aquatic and terrestrial vertebrates and the underlying mechanisms.

Our prediction that freshwater and marine invertebrates would be significantly different for body %N, but not P in aquatic invertebrates was supported by our findings. Freshwater and saltwater pose fundamentally different physical and nutrient challenges to animals. These mediums have different densities, which affect both buoyancy and the force required to move through the medium (Alexander 1990). They also have different nutrient cycles with freshwater environments having stereotypically more allochthonous carbon sources and lower ionic content than saltwater habitats (Bell *et al*. 2004; Boros *et al*. 2015; Hobbie 1988). Unfortunately, due to the lack of records for marine vertebrates, our data only allowed us to examine differences between invertebrates in the two aquatic realms. We are not aware of previous studies that have specifically compared the elemental content of freshwater and marine invertebrates. Elemental differences between organisms inhabiting these environments could also be driven by phylogeny and could underlie physiological and developmental differences between these two distantly-related groups. Therefore, further research will be necessary to disentangle the drivers underlying elemental differences between these two groups.

## Conclusions

Based on the first comprehensive comparison of terrestrial and aquatic animals across vertebrate and invertebrate taxa, our study sheds light on the role of body size, realm, and taxonomy in driving variation in the elemental content of animals. Neither body size nor realm alone explained the large observed variation in %N, %P, and N:P across animal species, with most of the variation being explained by the taxonomy. These findings starkly contrast with other biological attributes such as metabolic rate, which scale closely with body size across taxonomic groups (Kleiber 1932; Savage *et al*. 2004, 2007). Understanding animal elemental stoichiometry and therefore their nutrient demands, can help predict shifts in community content and biomass distribution across size classes as a response to changes in the availability of nutrients (Elser *et al*. 1996; Filipiak & Filipiak 2022; Hall 2009). In such analyses, stoichiometry needs to be considered explicitly, in addition to other traits such as body mass and metabolic rate. While body size was a poor predictor of the elemental content across species of vertebrates and invertebrates, we did find strong intraspecific patterns in the scaling of elements, consistent with tissue-scaling relationships that hold broadly across large groups of animals. These patterns could provide valuable information to help predict the responses of individual species to ongoing and future changes in major biogeochemical cycles driven by anthropogenic activities (Battye *et al*. 2017; Peñuelas *et al*. 2013; Tipping *et al*. 2014). Further research assessing how these scaling relationships within and among taxonomic groups, can help uncover the evolutionary and ecological drivers of stoichiometric diversity, specifically for terrestrial vertebrates and non-arthropod invertebrates. Our analyses on animal stoichiometry, using the largest and most comprehensive database to date, failed to find major size scaling relationships observed in studies examining smaller datasets of closely related animals. Instead, taxonomic identity, which is a proxy for physiological and developmental traits of animals, seems to be much more important at predicting the elemental content of individual organisms.

## Acknowledgements

We acknowledge funding of iDiv via the German Research Foundation (DFG FZT 118, 202548816), specifically funding through sDiv, the Synthesis Centre of iDiv. We especially thank Marten Winter and the sDiv team for their support in organizing the sBiomaps working group, which led to the compilation of the StoichLife database. This research was partially supported by a grant to ALG. from the National Science Foundation (NSF) DEB-1754326. MF was supported by Jagiellonian University, Faculty of Biology (N18/DBS/000003). NJG was supported by the US National Science Foundation (NSF) grant 2019470. AP was funded by the Deutsche Forschungsgemeinschaft (DFG, German Research Foundation) – Projektnummer 493345801. AT was supported by the Russian Science Foundation, project No. 22–14-00363. GQR was supported by Sao Paulo Research Foundation (FAPESP, grants 2019/08474-8 and 2022/10765-3), CNPq-Brazil productivity Grant, and funding from the Royal Society, Newton Advanced Fellowship (Grant No. NAF/R2/180791).

## Author contributions

MPN and ALG developed the idea for this paper. MPN led the statistical analyses and visualization, and wrote the first draft of the manuscript with input from ALG, OD, JM, and NJG. ALG and OD worked in database construction and curation with the help of sBiomaps participants KA, UB, MF, HH, SH, MJ, SL, MPN, REO, GP, RP, MS, AR, ES, JS, and EZ. ALG and OD obtained the iDiv funding that supported the sBiomaps working group. All other authors contributed data. All authors provided feedback on multiple drafts of this manuscript, and helped edit the manuscript.

