## Supplementary Figures for "Body size is a better predictor of intra- than interspecific variation of animal stoichiometry across realms"

**Appendix**

| Vertebrate | | | | | | | | |
| --- | --- | --- | --- | --- | --- | --- | --- | --- |
|  |  | Intercept | Slope - All | Slope - Aquatic | | Slope - Terrestrial | Marginal R^2^ | Conditional R^2^ |
| %N | Mean | **1.006***** | -0.008 | -0.010 | | -0.005 | 0.016 | 0.593 |
|  | SE | 0.026 | 0.013 | 0.006 | | 0.027 |  |  |
| %P | Mean | 0.315 | 0.021 | < -0.001 | | < 0.001 | 0.031 | 0.844 |
|  | SE | 0.229 | 0.078 | 0.020 | | 0.182 |  |  |
| Invertebrate | | | | | | | | |
|  |  | Intercept | Slope - All | Slope - Freshwater | Slope - Marine | Slope - Terrestrial | Marginal R^2^ | Conditional R^2^ |
| %N | Mean | **0.784***** | -0.007 | 0.006 | **-0.032**** | 0.003 | 0.010 | 0.904 |
|  | SE | 0.083 | 0.004 | 0.006 | 0.010 | 0.003 |  |  |
| %P | Mean | **-0.374*** | **-0.028**** | -0.020 | -0.017 | **-0.049***** | 0.015 | 0.793 |
|  | SE | 0.127 | 0.010 | 0.013 | 0.024 | 0.013 |  |  |

**Extended Data Table 1 |** Output of LMMs for log_10_-log_10_ transformed %N and %P of vertebrates and invertebrates for all animals (interspecific scaling). Slopes and intercepts of different realms using estimated marginal means. Predictions are on the population level (fixed effects). The intercept column represents the marginal mean of both groups at a body mass of 1. The column “Slope - All'' represents the marginal mean slope across realm levels, so that its magnitude is showing the average effect. The slopes of each realm are also shown. Mean and standard error (SE) values are given for all fixed-effect terms as marginal and fixed plus random-effect terms as conditional R^2^ values, for the full model. Significant values are indicated by bold font and asterisks (*** = *P* <0.001 ** = *P* < 0.01 * = *P* < 0.05).

**Extended Data Table 2 |** Full list of scaling relationships for each species analyzed. For each species the element measured (N, P, N:P), group (vertebrate, or invertebrate), specific habitat (freshwater, marine terrestrial), complete taxonomy (class, order, family, species), number of samples (n), scaling slope, R^2^ and *P*-value. Significant scaling relationships (*P* < 0.05) are shown in bold.

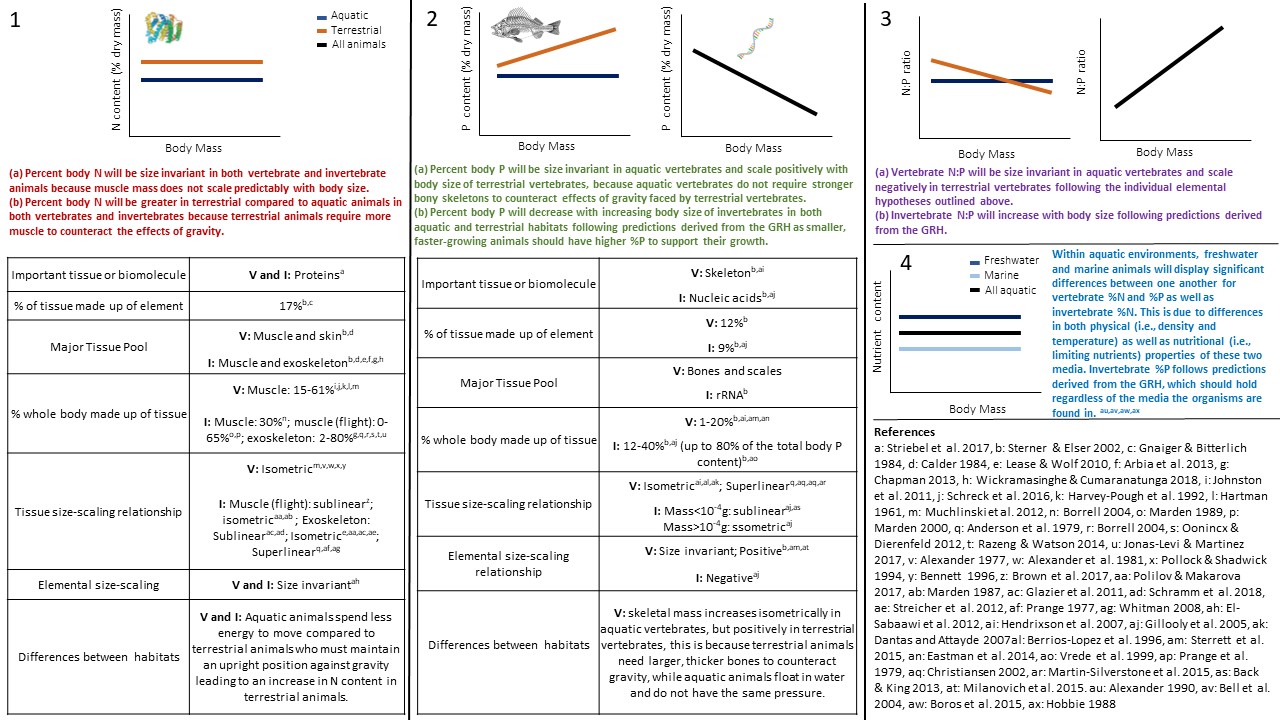

**Extended Data Figure 1 | Rationale and Evidence.** Rationale and empirical evidence on the size scaling of important body tissues for %N (**1**), %P (**2**) and N:P (**3**), as well as the difference in these scaling relationships in marine and freshwater realms (**4**), in vertebrates (V) and invertebrates (I), and derived predictions on the elemental content scaling with body mass.

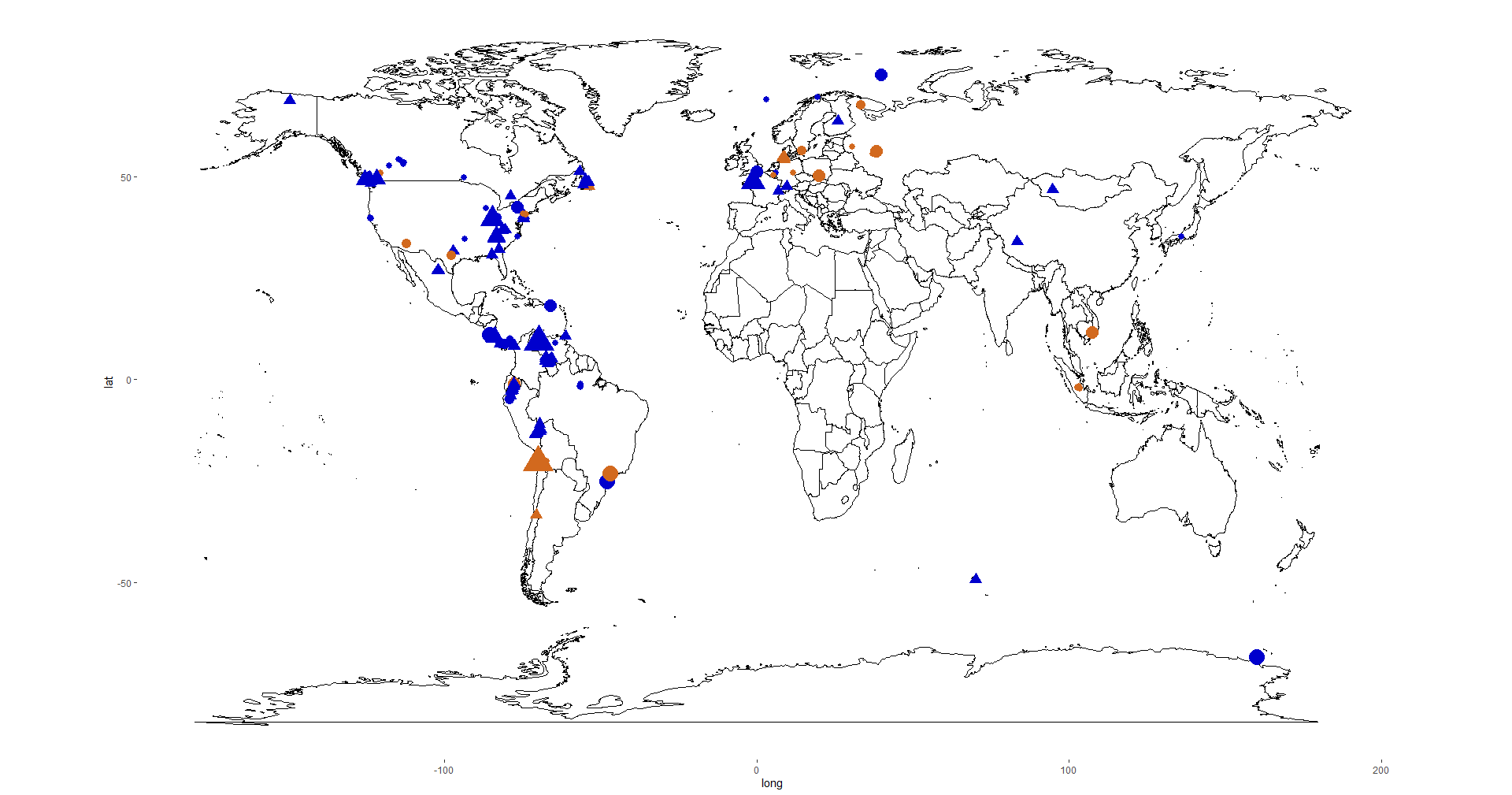

**Extended Data Figure 2 | Sample Map.** Map showing the sampling locations of animals included in the analysis. This data is a subset of StoichLife, restricting the database to only animals with elemental and body size measurements. Specimens were collected from 401 sampling locations (219 aquatic in blue; 182 terrestrial in brown). Vertebrates are indicated with a triangle, while invertebrates are shown with a circle. The size of the symbol scales with the number of individuals collected at the sampling locations.

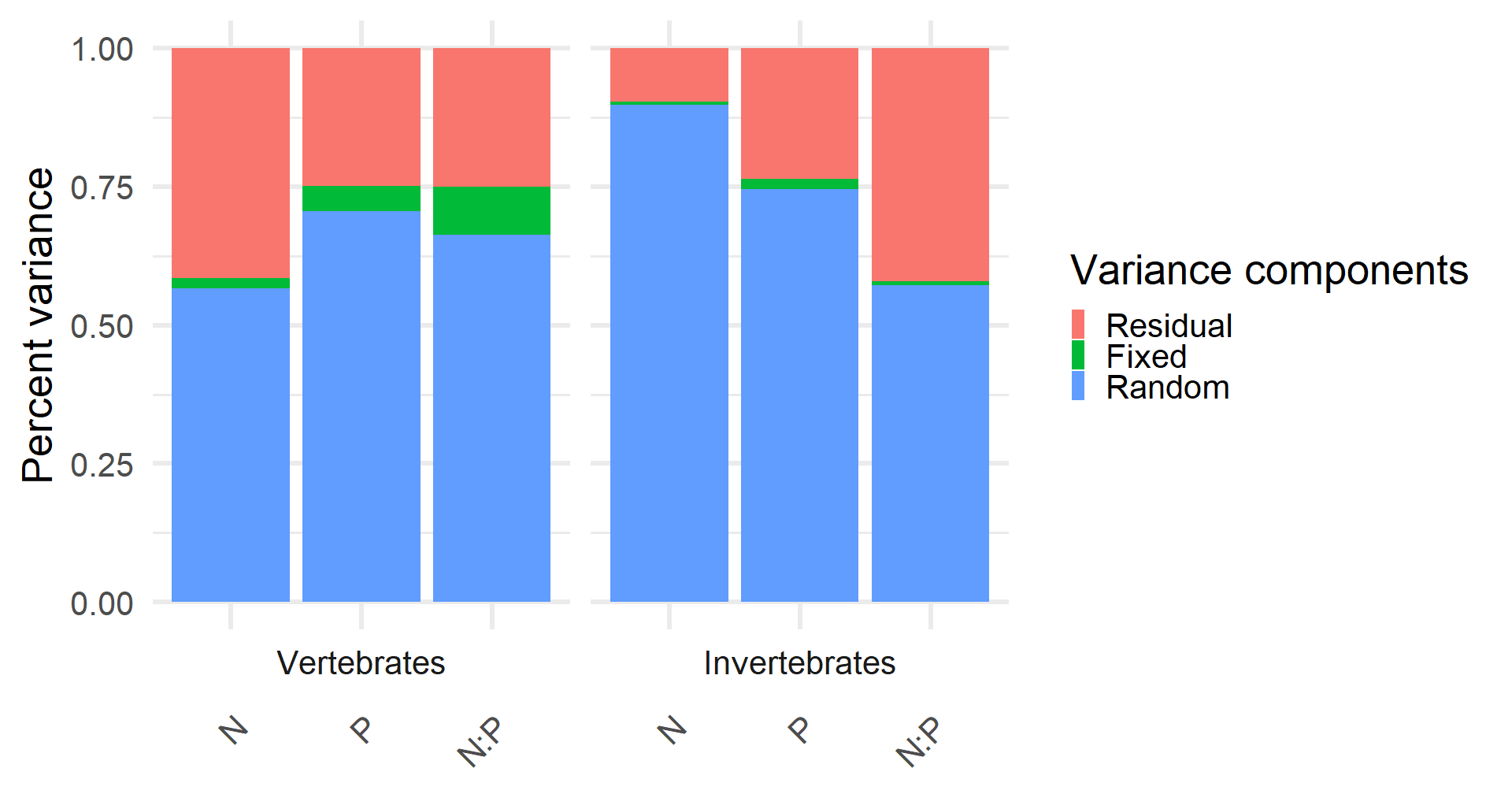

**Extended Data Figure 3 | Variance components.** Variance explained by the fixed and random effects (taxonomic group) of the LMMs for vertebrates and invertebrates and each element and ratio. For vertebrates, family was used for %N while class was used for %P and N:P. For invertebrates, phylum was used for %N and %P, and family for the N:P ratio.

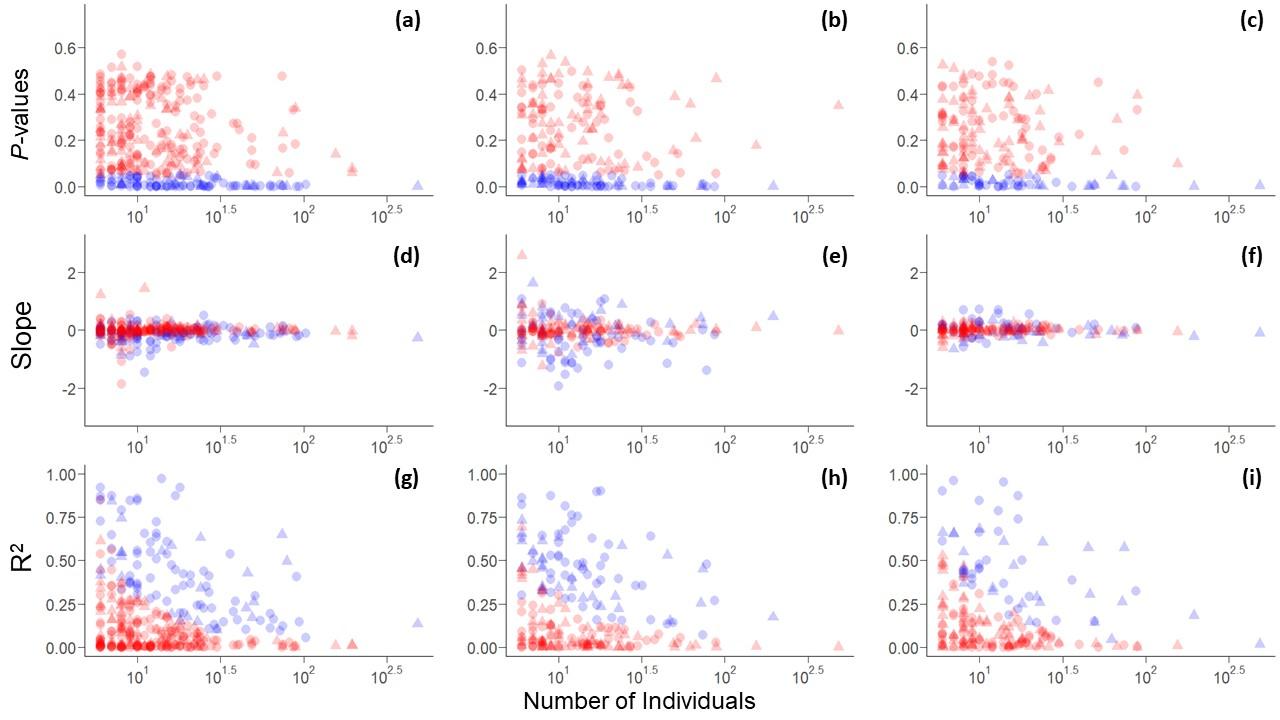

**Extended Data Figure 4 |** **Sample size relationships.** Intraspecific models for each species plotted as a function of the number of observations (n ranges from 6 to 481) for each species for %N (**a, d, g**), %P (**b, e, h**) and N:P (**c, f, i**) for *P*-values (top), slopes (middle), and R^2^ values (bottom). Relationships scored as significant (*P* < 0.05) are shown in blue while non-significant relationships are shown in red. Each point represents a single species with vertebrates represented as triangles and invertebrates as circles.
